# Deciphering the major metabolic pathways associated with aluminum tolerance in popcorn roots using label-free quantitative proteomics

**DOI:** 10.1101/2021.08.22.455814

**Authors:** Vitor Batista Pinto, Vinícius Costa Almeida, Ítalo Antunes Pereira Lima, Ellen de Moura Vale, Wagner Luiz Araújo, Vanildo Silveira, José Marcelo Soriano Viana

## Abstract

Aluminum toxicity is one of the most important abiotic stresses that affect crop production worldwide. The soluble form (Al^3+^) inhibits root growth by altering water and nutrients uptake, which also reduces plant growth and development. Under a long term Al^3+^ exposure, plants can activate several tolerance mechanisms, and to date, there are no reports of large-scale proteomic data of maize in response to this ion. To investigate the post-transcriptional regulation in response to Al toxicity, we performed a label-free quantitative proteomics for comparative analysis of two Al-contrasting popcorn inbred lines and an Al-tolerant commercial hybrid during 72 h under Al-stress. A total of 489 differentially accumulated proteins (DAPs) were identified in the Al-sensitive inbred line, 491 in the Al-tolerant inbred line, and 277 in the commercial hybrid. Among them, 120 DAPs were co-expressed in both Al tolerant genotypes. Bioinformatics analysis indicated that starch and sucrose metabolism, glycolysis/gluconeogenesis, and carbohydrate metabolism were significant biochemical processes regulated in response to Al toxicity. The up accumulation of sucrose synthase and the increase of sucrose content and starch degradation suggest that these components may enhance popcorn tolerance to Al stress. The up-accumulation of citrate synthase suggests a key role of this enzyme in the detoxification process in the Al-tolerant inbred line. The integration of transcriptomic and proteomic data indicated that the Al tolerance response presents a complex regulatory network into the transcription and translation dynamics of popcorn roots development.

## 1. Introduction

Aluminum (Al) toxicity occurs in approximately 30% of the world’s arable soils and in more than 50% of potentially arable land. From this total, approximately 60% is located in tropical and subtropical regions, negatively impacting the food supply chain. In acid soils, with pH values of 5 or below, the phytotoxic species Al^3+^ is solubilized in soil solution and becomes one of the most important abiotic stresses limiting crop production. The phytotoxic form Al^3+^ inhibits root growth, altering water and nutrients uptake, which also reduces plant growth and development (von Uexküll and Mutert 1995; Kochian et al. 2002, 2015).

The physiological aspects of Al resistance have been well understood. To cope with Al toxicity, plants developed multiple strategies including the mechanisms of Al-exclusion and detoxification. The exclusion mechanism prevents the Al entrance in the root apex via limiting the binding of the Al ion to the root cell wall, altering the selective permeability of the plasma membrane and releasing organic acids, phosphate, and phenolic compounds from the roots to attenuate Al-toxicity (Pellet et al. 1995; Kidd et al. 2001; Mattiello et al. 2010; Yang et al. 2011). The mechanism underlying internal Al tolerance involves the detoxification and sequestration into the vacuole forming a stable and non-phytotoxic complex (Delhaize and Ryan 1995; Ma 2005).

The well-characterized exclusion mechanism is dependent on organic acid (OA) exudation from the root apex (Ma et al. 2001; Kochian et al. 2004). Al exposure induces malate secretion in wheat (Delhaize et al. 1993), Arabidopsis (Hoekenga et al. 2006), and rapeseed (Ligaba et al. 2006); citrate secretion in sorghum (Magalhaes et al. 2007), barley (Furukawa et al. 2007), rice (Yokosho et al. 2011), wheat (Ryan et al. 2009; Tovkach et al. 2013), and maize (Maron et al. 2013), and oxalate secretion in tomato (Yang et al. 2011), buckwheat (Jian Feng Ma et al. 1997), and spinach (Jian et al. 2005).

Maize is widely grown on acid soils throughout the tropics and subtropics (Maron et al. 2013), and Al tolerance is an important trait in breeding programs in these regions. In Brazil, maize cultivation reached around 4,156.6 thousand hectare of planted area and the productivity of 6,338 kg ha^−1^ in the 2019/2020 harvest (CONAB 2020). In addition, popcorn (*Zea mays* var. *everta*) cultivation is in increasing expansion, attracting the attention of breeders to obtain populations and hybrids adapted to Brazilian conditions (Moterle et al. 2012).

Although many genes have been identified in controlling different Al-tolerance mechanisms in maize (Maron et al. 2010; Guimaraes et al. 2014; Matonyei et al. 2020), the physiological and cellular responses induced by Al-stress and the genetic components of Al-tolerance pathways are not well understood (Duressa et al. 2011). Thus, proteomic analysis provides insights into post-transcriptional regulation that controls cellular functions complementing the genomic analysis (Ralhan et al. 2008). This is the only possible way to identify the genetic and molecular components that underlie the Al-tolerant phenotype (Duressa et al. 2011).

Some research has been conducted with the analysis of the root proteome of plants under Al-stress (Yang et al. 2007; Zhen et al. 2007; Duressa et al. 2011; Wang et al. 2014; Oh et al. 2014), but its major limitation is to integrate these data with metabolite measurements to understand the biochemical pathways that are needed for the Al-tolerance response. In this study we identified differentially accumulated proteins responsive to the Al^3+^ toxicity in popcorn genotypes differing in Al tolerance, using mass spectrometry. To our knowledge, there are no reports to date, addressing proteins responsive to Al using the proteomics approach and its integration with metabolic measurements in *Zea mays* var. *everta*.

## 2. Material and methods

### 2.1. Plant material

Two Al-contrasting popcorn inbred lines were developed by the Popcorn Breeding Program at Universidade Federal de Viçosa (Viçosa, MG, Brazil). Previously, 18 inbred lines were evaluated for relative root growth (RRG), hematoxylin staining, Al content, and external morphology of roots, and the inbred line 11-133 was selected as the most Al-tolerant and the inbred line 11-60 was selected as the most Al-sensitive (Rahim et al. 2019). Moreover, the Al-tolerant commercial hybrid 2B710PW from Dow AgroSciences was included in the experiments.

### 2.2. Growth conditions

Initially, seeds were treated with fungicide (Captan-400®) and germinated at 25 °C + 1 °C in a growth chamber for 7 days. Seedlings with uniform growth were picked randomly and transferred to a nutritive solution with constant aeration to acclimated for 24 h. The nutrient solution composition was: 1 mM KCl, 1.5 mM NH_4_NO_3_, 1 mM CaCl_2_, 45 µM KH_2_PO_4_, 200 µM MgSO_4_, 500 µM Mg(NO_3_)_2_, 155 µM MgCl_2_, 11.8 µM MnCl_2_.4H_2_O, 33 µM H_3_BO_3_, 3.06 µM ZnSO_4_.7H_2_O, 0.8 µM CuSO_4_.5H_2_O, 1.07 µM Na_2_MoO_4_.H_2_O, and 77 µM Fe-EDTA (Magnavaca et al. 1987; Famoso et al. 2010). Then, the treatment group was submitted to aluminum stress with 160 μM Al^3+^ at pH 4.5 for 72 h. The seedlings were maintained in a growth chamber at 25°C with 12/12 h light/dark cycle.

### 2.3. Root growth measurements

After seven days, the seedlings were removed from the solutions, and the roots were scanned with EPSON LA2400 Perfection V700/V750 scanner equipped with additional light (transparency unit) at a resolution of 400 dpi. The WinRhizo software (Bouma et al. 2000) was used to measure the root growth parameters.

### 2.4. Aluminum concentration

The Al concentrations were quantified in total roots and root tips (0.5 cm) after 7 days in nutritive solution (Silva et al. 2020). The samples were analyzed in an inductively coupled plasma optical emission spectroscopy (ICP-OES, Perkin-Elmer Optima 3000XL, Maryland, USA).

### 2.5. Metabolite quantification

Total roots were harvested and immediately frozen in liquid nitrogen, then stored at -80 °C until use. The metabolite extraction was performed with an ethanol gradient (98%, 80% and 50%) in cycles of 80 °C. Subsequently, glucose, sucrose, and starch, concentrations were measured as described by Fernie et al. (2001) and Cross et al. (2006). Malate and fumarate contents were determined according to Nunes-Nesi et al. (2007).

### 2.6. Proteomic analysis

Total protein extraction was performed according to Damerval et al. (1986), with modifications. Total roots from each treatment were sampled at 72 h after Al-stress. Samples (300 mg of fresh material each, in three biological replicates) were first macerated to a fine powder with liquid nitrogen, resuspended in 1 mL of extraction buffer containing 10% (w/v) trichloroacetic acid (TCA; Sigma-Aldrich; St. Louis, USA) in acetone (Merck; Darmstadt, Germany) with 20 mM dithiothreitol (DTT; GE Healthcare; Piscataway, USA), vortexed for 5 min at 4 °C, and kept at – 20 °C for 1 h and then centrifuged at 16000 g for 30 min at 4 °C. The resulting pellets were washed three times with cold acetone containing 20 mM DTT and subsequently air dried. The pellets were resuspended in buffer containing 7 M urea (GE Healthcare), 2 M thiourea (GE Healthcare), 1% DTT, 1 mM phenylmethylsulfonide (PMSF; Sigma-Aldrich), complete protease inhibitor cocktail (Roche Diagnostics, Mannheim, Germany) and 2% Triton X-100 (Sigma-Aldrich), vortexed for 30 min, and then centrifuged at 16000 g for 20 min at 4 °C. The supernatants were collected, and the protein concentrations were determined using a 2-D Quant Kit (GE Healthcare; Piscataway, USA).

The proteins samples were precipitated using a methanol/chloroform methodology to remove any interferents from the samples (Nanjo et al. 2012). The samples were then resuspended in a solution consisting of 7 M urea and 2 M thiourea, and tryptic protein digestion (1:100 enzyme:protein, V5111, Promega, Madison, USA) was performed using Microcon-30 kDa filter units (Millipore) with the filter-aided sample preparation (FASP) methodology (Reis et al. 2021). The resulting peptides were quantified with the A205 nm protein and peptide methodology using a NanoDrop 2000c spectrophotometer (Thermo Fisher Scientific, Waltham, USA).

The samples were then injected into a nanoAcquity UPLC mass spectrometer connected to a Q-TOF SYNAPT G2-Si instrument (Waters, Manchester, UK). The runs consisted of three biological replicates of 1 μg of peptide samples. The spectra processing and database search were performed using the ProteinLynx Global SERVER (PLGS) software v.3.02 (Waters) and the *Zea mays* protein databank (ID: UP000007305) available on UniProtKB (www.uniprot.org). The label-free quantification analyses were performed using ISOQuant software v.1.7 (Distler et al. 2014). To ensure the quality of the results after data processing, only proteins present in the three runs were accepted for differential abundance analysis. Proteins were deemed up-accumulated if the log2 value of the fold change (FC) was greater than 0.60 and down-accumulated if the log2 value of the FC was less than − 0.60, according to Student’s t test (two-tailed, P < 0.05) and after P value correction with the Benjamini-Hochberg test. The functional enrichment analysis was performed using OmicsBox software 1.2.4 (https://www.biobam.com/omicsbox) and Metascape (Zhou et al. 2019) (http://metascape.org).

The mass spectrometry proteomics data, the complete description step-by-step of the proteomic methods and setting parameters, and the complete list of identified proteins from proteomics analysis have been deposited in the ProteomeXchange Consortium (Deutsch et al. 2017) via the PRIDE (Perez-Riverol et al. 2019) partner repository, with the dataset identifier PXD027588.

## 3. Results

### 3.1. Root growth and aluminum content

The root growth analysis was performed after seven days under Al-stress. The phenotypic differences between the tested genotypes can be visualized in Figure 1A. In the three genotypes was detected a decrease in the total root length (TRL) when growing under Al-stress (Figure 1B). The same was visualized on the root surface area (RSA), except for the Al-tolerant inbred line that not differs statistically from the control condition (Figure 1C). In contrast, the Al-stress was responsible to increase the root diameter (RAD) of all the genotypes (Figure 1D). The number of roots decreased in the two Al-tolerant genotypes when compared to the control condition (Figure 1E). For Al content, it was visualized that the Al-tolerant inbred line accumulated less Al in the root tip and in total root when compared to the other tested genotypes (Figure 1F).

**Figure 1.**
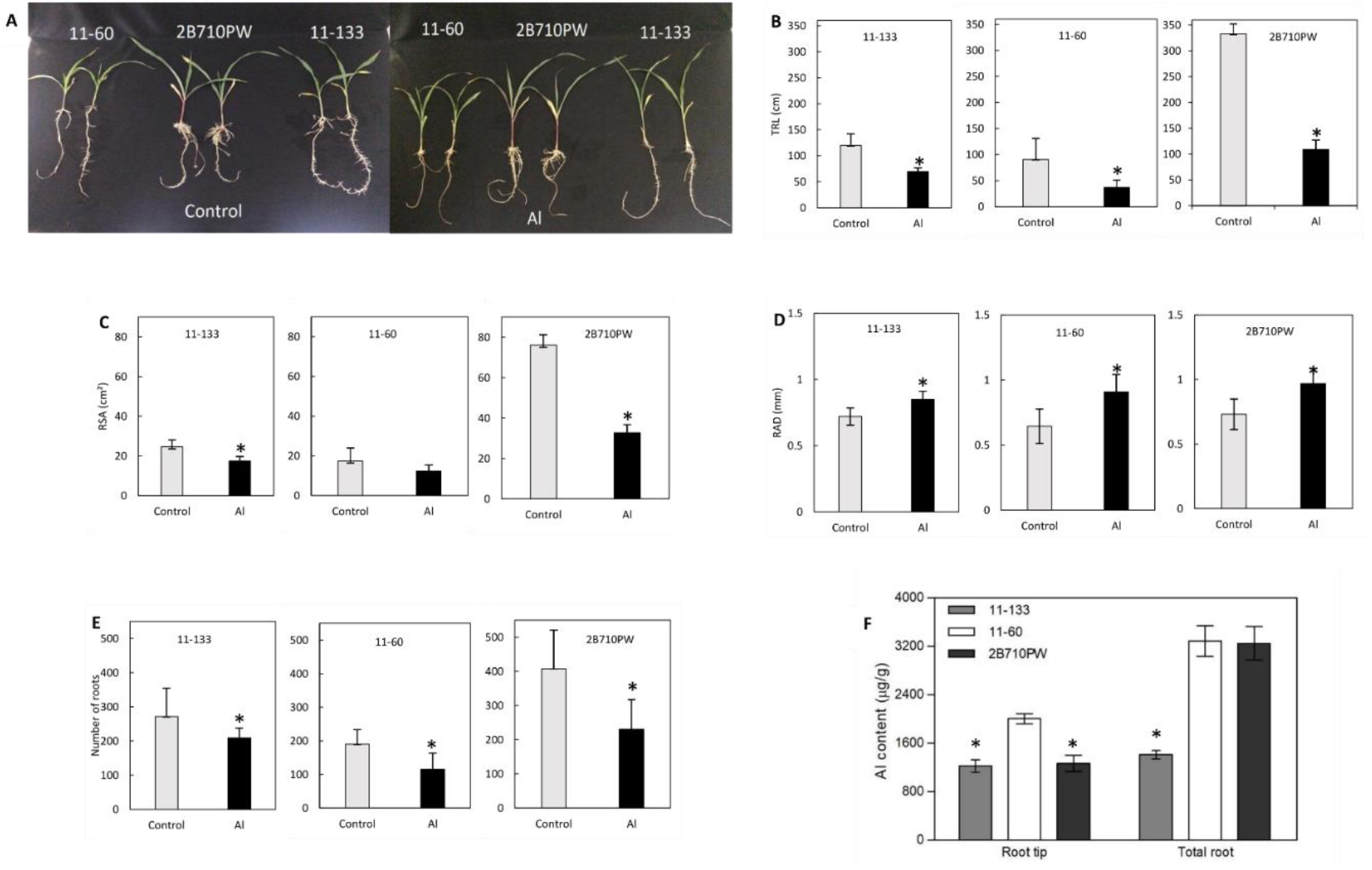
Growth measurements and aluminum content. **(A)** Phenotype of the three tested genotypes after seven days under control and Al-stress. **(B – E)** Root growth measurements. Values followed by asterisks are statistically different from the control by the Student’s t-test at 5%. Values are means ± SE (n=15). Total root growth (TRG); root surface area (RSA); root diameter (RAD). **(F)** Al content in the root tip and in total root after seven days under Al-stress. Values followed by asterisks are statistically different from the control by the Student’s t test at 5%. Values are means ± SE (n=5).

### 3.2. Proteomic profile

A total of 489 DAPs were detected in the Al-sensitive inbred line, 491 in the Al-tolerant inbred line, and 277 in the commercial hybrid (Figure 2A; Suppl. Table S1). From this total, were detected 71; 31; and 38 unique proteins in the control condition, and 57; 77; and 49 unique proteins in Al-stress condition in the Al-sensitive, Al-tolerant, and commercial hybrid, respectively (Suppl. Table S1). A total of 120 DAPs were co-expressed in the genotypes resistant to Al ion (Figure 2B). The top 20 significant enriched GO functional groups of DAPs (P < 0.05) was obtained with Metascape. The DAPs were classified into three ontologies (Figure 2C). The most enriched functional group in the three contrasts was response to cadmium ion (GO:0046686), followed by cell wall (GO:0005618) in both inbred lines, and cytosolic part (GO:0044445) in the genotypes tolerant to Al toxicity.

**Figure 2.**
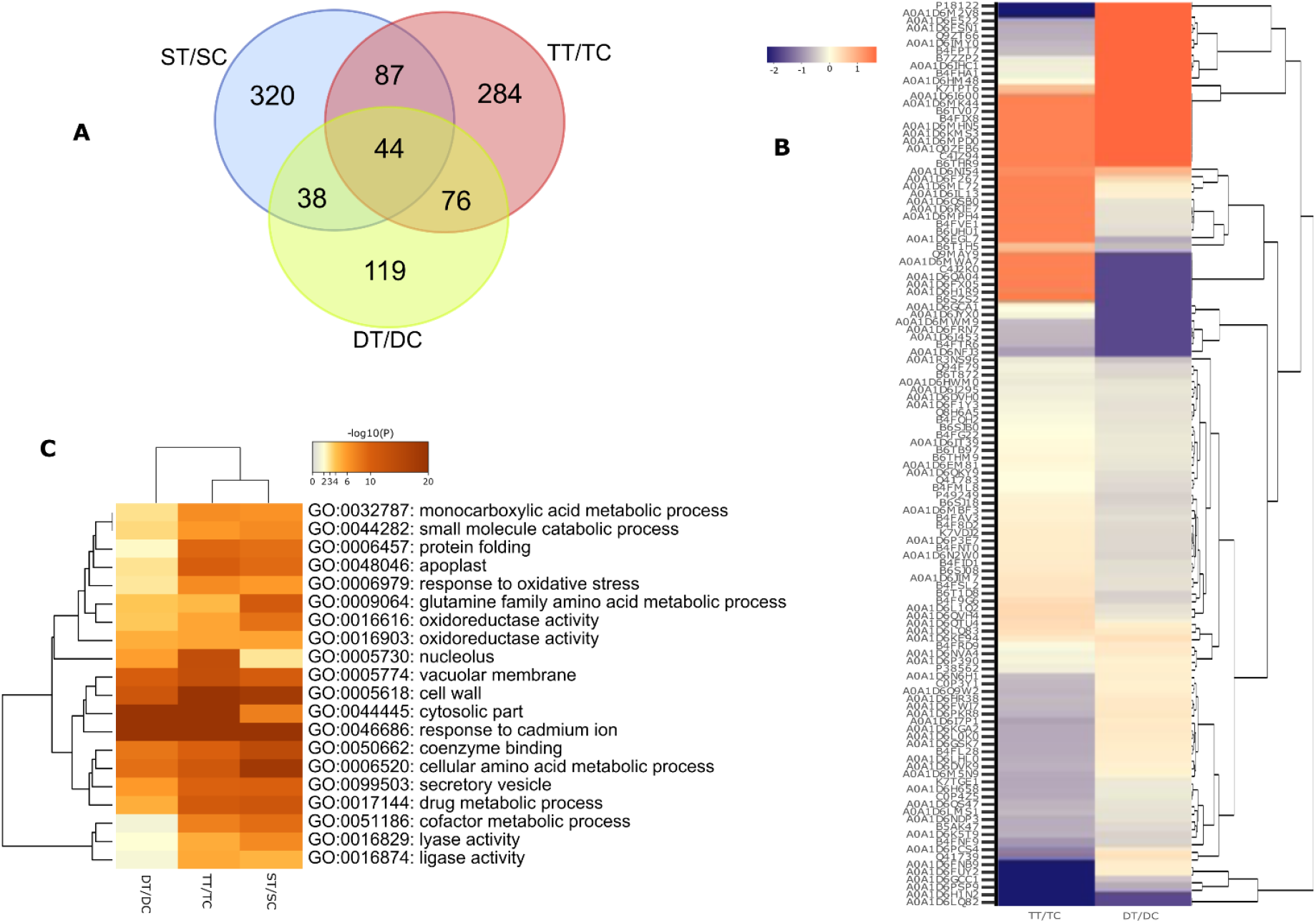
Proteomic analysis of popcorn roots under 72 hours of Al-stress. **(A)** Venn diagram shows the overlap differentially accumulated proteins. (ST/SC: contrast between Al-sensitive inbred line under Al-stress/control; TT/TC: contrast between the Al-tolerant inbred line under Al-stress/control; DT/DC: contrast between commercial hybrid under Al-stress/control). **(B)** GO enrichment of top 20 GO terms of DAPs using Metascape *Arabidopsis thaliana* database. **(C)** Heatmap of UPGMA clustering of Al co-regulated proteins between the Al-tolerant inbred line and commercial hybrid.

The DAPs were classified and grouped according to their primary biological activities, as defined by the Clusters of Orthologous Groups (COG). The DAPs from the genotypes were classified into 21 COG categories (Figure 3A). The most representative categories in the three tested genotypes were “post-translational modification, protein turnover, and chaperones” (group O), which retrieved the major number of proteins in the Al-sensitive inbred line, and “function unknown” (group S). In the two genotypes tolerant to the Al ion, the category with the major number of proteins was “translation, ribosomal structure and biogenesis” (group J). In the commercial hybrid, no proteins were categorized in the category “chromatin structure and dynamics” (group B).

**Figure 3.**
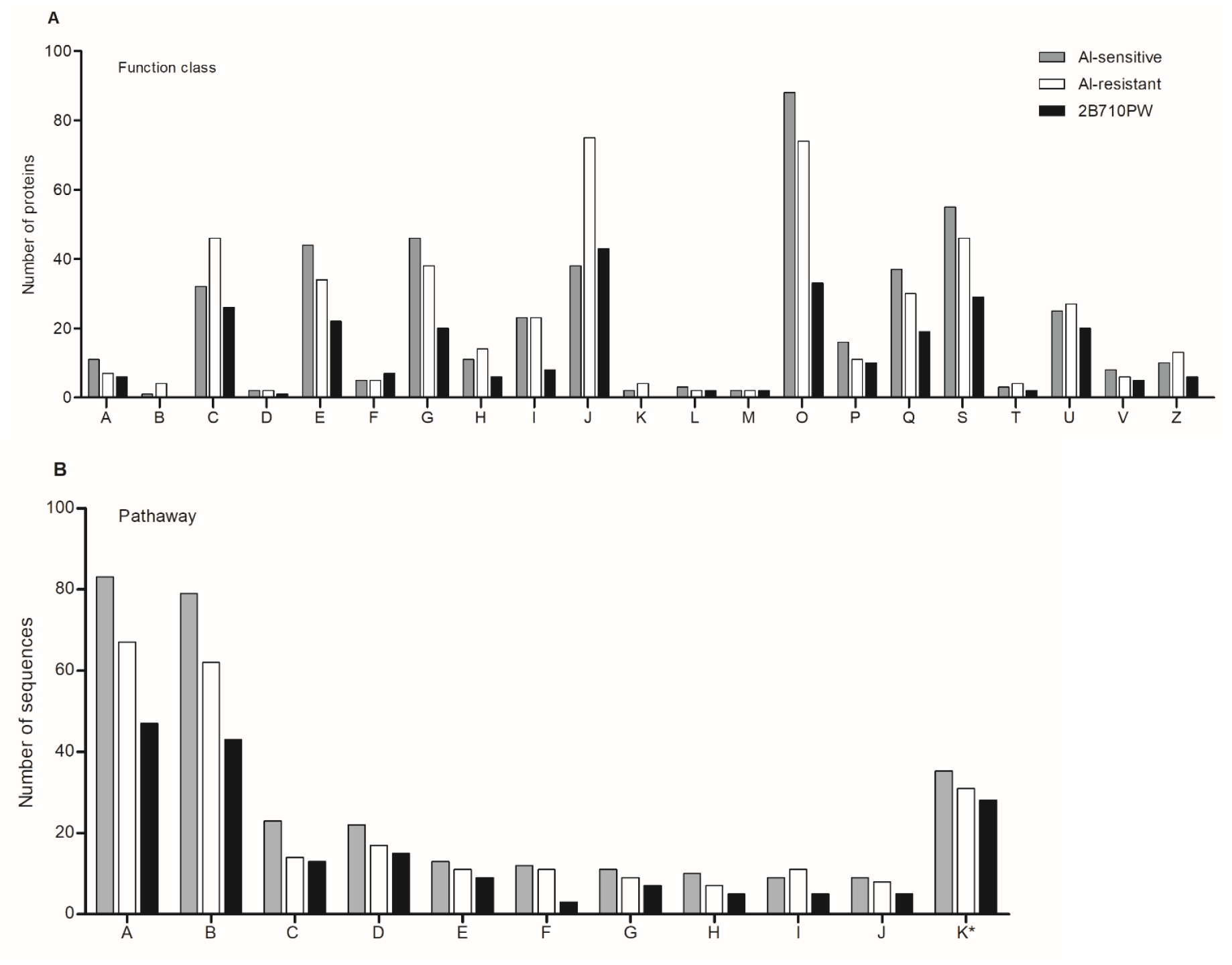
COG classification and KEGG pathway analysis of DAPs under Al stress. **(A)** Functional classes produced by COG. The abbreviations in the figure (A) are as follows: A, RNA processing and modification; B, Chromatin structure and dynamics; C, Energy production and conversion; E, Amino acid transport and metabolism, Coenzyme transport; F, Nucleotide transport and metabolism; G, Carbohydrate transport and metabolism; I, Lipid transport and metabolism; J, Translation, ribosomal structure and biogenesis; K, Transcription; L, Transcription, Replication, recombination and repair; M, Cell wall/membrane/envelope biogenesis; O, Post-translational modification, protein turnover, and chaperones; P, Inorganic ion transport and metabolism; Q, Secondary metabolites biosynthesis, transport, and catabolism; S, Function unknown; T, Signal transduction mechanisms; U, Intracellular trafficking, secretion, and vesicular transport; V, Defense mechanisms; Z, Cytoskeleton. **(B)** KEGG pathways enrichment. The abbreviations in the figure (B) are as follows: A, Purine metabolism; B, Thiamine metabolism; C, Glycolysis/Gluconeogenesis; D, Pyruvate metabolism; E, Alanine, aspartate and glutamate metabolism; F, Glutathione metabolism; G, Citrate cycle (TCA cycle); H, Carbon fixation in photosynthetic organisms; I, Starch and sucrose metabolism; J, Amino sugar and nucleotide sugar metabolism; K, Other pathways. * The number of DAPs were divided by ten.

A KEGG pathway analysis was performed to investigate the biological function of the DAPs. A total of 624; 527; and 433 sequences from the inbred lines 11-60 and 11-133, and the commercial hybrid, respectively, were mapped to several pathways. The most represented pathway was “purine metabolism”, followed by “thiamine metabolism”, and “glycolysis/gluconeogenesis” (Figure 3B). The commercial hybrid presented the lowest number of proteins mapped into the KEGG pathways.

### 3.3. Metabolite quantification

Malate and fumarate increased in the three genotypes after 72 h in the roots under stress when compared to the control. In stress condition, the shoots of the commercial hybrid accumulated the highest average levels of malate in comparison with the other inbred lines (Table 1).

**Table 1.**
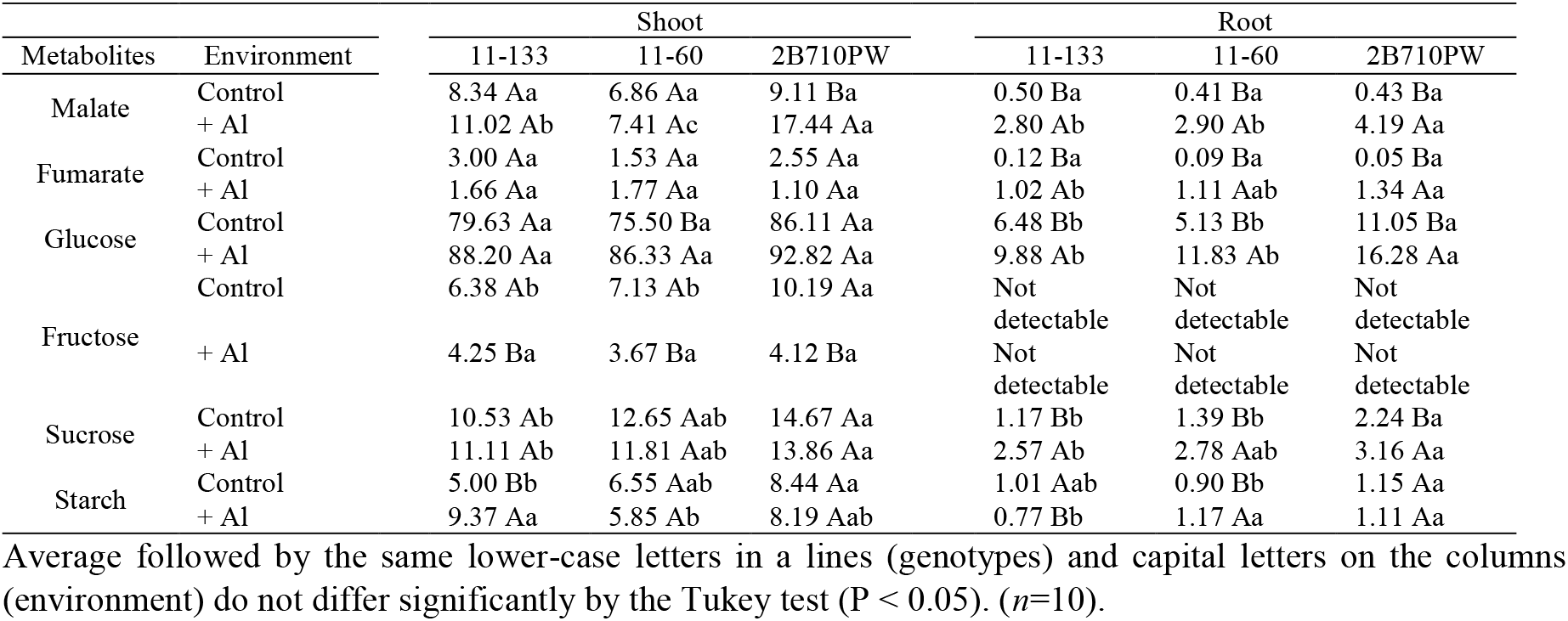
Primary metabolites in popcorn genotypes under aluminum (Al) exposure.

The glucose content increased in roots of plants under stress, presenting high average values in the commercial hybrid (Table 1). It was not detected fructose in the roots of all the tested genotypes and a decrease of its levels was observed in the shoots of plants under stress (Table 1). It was observed an increase of sucrose in the roots of all the genotypes under Al-stress (Table 1). Besides that, the sucrose content gradually increased in the roots and shoots in the Al-tolerant inbred line over the days under Al-stress (Suppl. Figure S1). In the same inbred line was detected a decrease of starch content in the roots and its consequent increase in the shoots under Al-stress (Table 1). This inbred line presented the lowest levels of starch in the roots in comparison with the other genotypes under stress, and the opposite was observed in the shoots (Table 1).

### 3.4. Transcriptomic and proteomic data integration

Integrating RNA sequencing data of the Al-tolerant inbred line (11-133) (NCBI SRA repository, accession code: PRJNA508768) with the proteomics analysis, both in the same conditions, were detected only 20 components regulated at transcriptional and post-transcriptional levels (Figure 4A; Suppl. Table S2). KEGG enrichment analysis of differentially expressed genes (DEGs) and DAPs showed that the Al-stress regulated players involved in bioenergetic metabolism mostly at the post-transcriptional level (Figure 4B).

**Figure 4.**
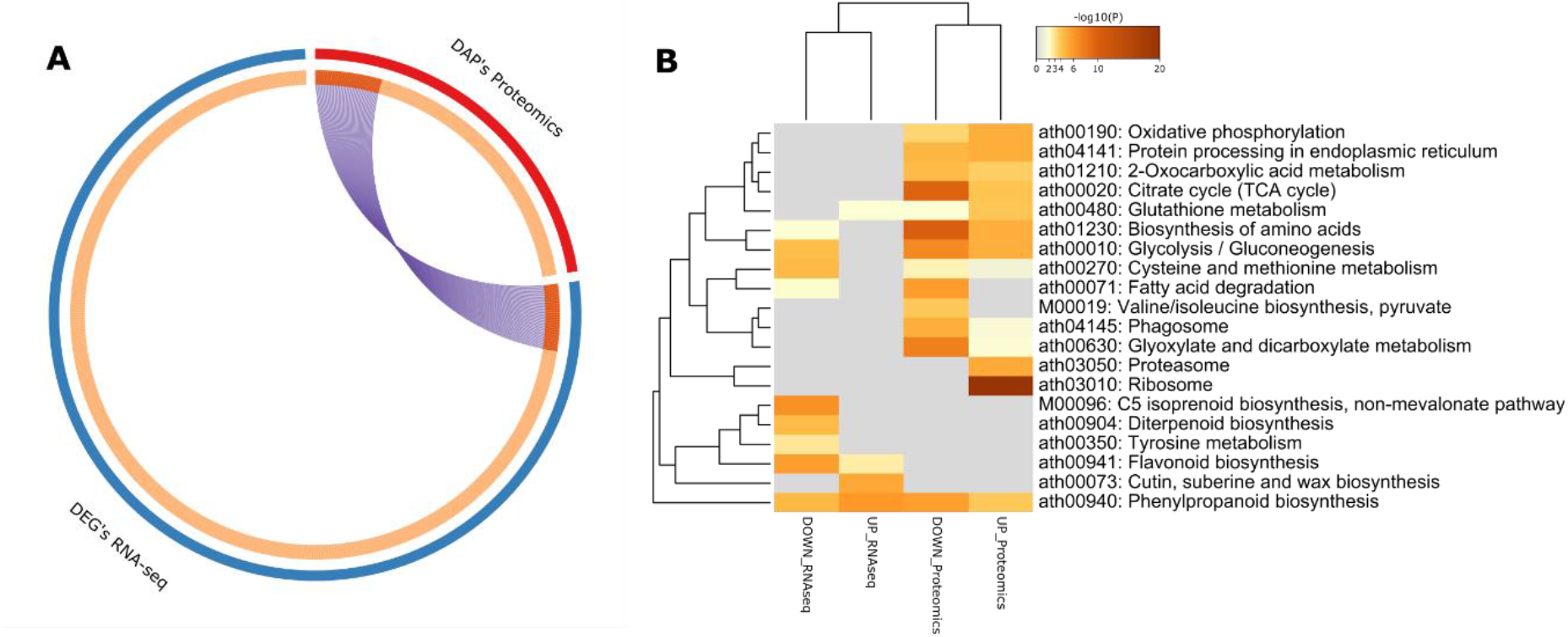
Integrative analysis of transcriptomic and proteomic data using Metascape *Arabidopsis thaliana* database. **(A)** Overlap between differentially expressed genes (DEGs) detected by RNA-seq and DAPs detected by label-free proteomics. The inner circle represents gene lists, where hits are arranged along the arc. Genes that hit both lists are colored in dark orange, and genes unique to a list are shown in light orange. **(B)** Heatmap of the top 20 cluster of KEEG enrichment pathways of DEGs and DAPs.

## 4. Discussion

### 4.1. Al stress affects root growth

We demonstrated that the Al-tolerant inbred accumulated fewer concentrations of Al in roots and root tips, not affecting the root surface area, but at the same time, a low concentration of Al is able to limit the total root length and the number of roots, and also increases the root diameter (Figure 1B-F). According to our data, the same Al-tolerant genotypes exhibited a higher total root growth comparing to the Al-sensitive inbred line (Rahim et al. 2019). Besides that, the degree of Al tolerance in maize is negatively correlated with the Al content in root tip once that an Al exclusion mechanism in the tolerant genotypes results in less Al accumulation in the tips of the roots than in sensitive genotypes (Piñeros et al. 2002).

### 4.2. Proteins regulated by Al stress

To track the proteomic changes of popcorn roots in a long-term response to Al toxicity, we performed a robust shotgun label-free proteomic analysis. From the total of proteins detected, 19.98%, 19.89%, and 11.27% were DAPs in Al-tolerant and Al-sensitive inbred lines, and in the commercial hybrid, respectively.

Comparing the both Al-tolerant genotypes, 120 DAPs were shared between them with different accumulation patterns (Figure 2B). In addition to the proteins discussed below, we highlight some co-expressed proteins that may contribute to the Al tolerance response.

A cysteine synthase mitochondrial (A0A1D6MPD0) was regulated only in Al-stress condition (Suppl. Table S1). Increased accumulation of this protein can accelerate the biosynthesis of cysteine, a precursor for the synthesis of several Al-tolerance related metabolites (Zheng et al. 2014) and it has been identified as up-regulated in roots of soybean (Duressa et al. 2011) and rice (Yang et al. 2007).

The Aluminum-induced protein (AIP) (A0A1D6IT39) and ABC transporter (A0A1D6I295) had their accumulation increased in the Al-tolerant inbred line but decreased in the commercial hybrid (Suppl. Table S1). The overexpression of AIP gene has been reported enhancing the Al tolerance in terms of root growth in Arabidopsis (Jang et al. 2014). ABC transporters are already known for their role in responses to environmental stress, mainly in Al detoxification in crops (Lane et al. 2016). This is one of the proteins that should be selected as biomarker to Al-tolerance in popcorn genotypes. Furthermore, differential proteins that responded to Al in some metabolic pathways are discussed below.

### 4.3. Protein synthesis, degradation and processing

The cells continuously adjust synthesis and degradation in order to maintain protein homeostasis in adverse environmental conditions (Hinkson and Elias 2011). The number of specific ribosomal proteins up-accumulated in our data indicates that the roots of the Al-tolerant inbred line detect the ion Al^3+^ and start to respond to it by increasing protein synthesis. In previous microarray analysis, the increased demand for specific ribosomal components was detected during Arabidopsis Al exposure (Kumari et al. 2008). The majority of elongation factors were down-accumulated in the Al-sensitive inbred line, contrasting with the up-accumulated in the Al-tolerant genotypes. The differential regulation in the components of the translation machinery indicates that the Al-tolerant genotypes have a distinct control of the protein synthesis in response to aluminum stress. Together, these data suggest that protein synthesis is strongly regulated in the Al-tolerant genotypes to survive Al stress.

Proteasomes were identified in the three genotypes. These data are according to those found in soybean (Duressa et al. 2011), tomato (Zhou et al. 2009), and rice (Yang et al. 2013), indicating that the protein degradation may be associated with the root growth inhibition mediated by Al toxicity.

Only three heat shock proteins (HSPs) were accumulated in the commercial hybrid, contrasting with 17 and 11 in the Al-sensitive and Al-tolerant inbred lines, respectively. However, chaperones and chaperonins were up-accumulated only in the Al-tolerant genotypes, while the majority were down-regulated or not detected in the Al-sensitive inbred line. Besides that, a T-complex protein 1 (A0A1D6MPH4) was regulated only in the Al-tolerant inbred line after Al^3+^ treatment. Several chaperones and HSPs were found differentially regulated in soybean (Duressa et al. 2011). In rice, a T-complex protein 1 had its accumulation increased in the Al-tolerant cultivar in contrast to the Al-sensitive (Wang et al. 2014). Based on this, the Al toxicity increases the activity of these proteins, consequently increasing the protein synthesis and turnover to survive in this condition.

### 4.4. Antioxidant system related proteins

In some plants, the Al toxicity might be responsible to the generation of reactive oxygen species (ROS) in mitochondria, chloroplast, and peroxisome (Kochian et al. 2005). In general, up-accumulated peroxidases were detected only in the Al-tolerant genotypes. Peroxidases were identified in root proteome of rice (Wang et al. 2014) and wheat (Oh et al. 2014), and in Arabidopsis the overexpression of AtPrx64 may improve tolerance to Al (Wu et al. 2017). The Al-stress increased the regulation of catalase (B6UHU1), catalase isozyme (CATA1), and thiamine thiazole synthase 1 (THI41) at least in one Al-tolerant genotype. Superoxide dismutase (B4F925) was detected in the commercial hybrid and none of the inbred lines, according to Pinto (Pinto 2019).

### 4.5. Starch and sucrose metabolism

Sucrose participates in plant response to abiotic stresses (Trouverie et al. 2003; Gupta and Kaur 2005) and takes part in the biosynthesis of starch, cellulose, and protein (Liu et al. 2018). The sucrose synthase (A0A1D6QA04) must be one of the key proteins against the Al toxicity in popcorn roots. This protein was identified in the Al-tolerant inbred line roots when exposed to Al-stress, and only in the control condition in both Al-sensitive and commercial hybrid (Suppl. Table S1).

Sucrose synthase was also induced in proteomics studies in the roots of rice (Wang et al. 2014). In cotton seedlings, the sucrose content was increased for utilization in cell wall formation under Al-stress (Huck 1972). The sucrose content was also increased in the roots in all genotypes in the Al-treatment but did not differ statistically in shoots (Table 1). Confirming our hypothesis that sucrose may be a key player in popcorn under Al-stress, as its content gradually increased in the roots and shoots in the Al-tolerant inbred line over the days (Suppl. Figure S1).

Adverse environmental factors can induce differential effects of source-sink on metabolism, leading to the differential expression of various proteins related to sucrose and carbohydrate metabolism (Koch et al. 1992; Roitsch et al. 1995). This way, sucrose can be either metabolized or transported to the vacuole, as its regulation is necessary for plant development and stress responses (Wind et al. 2010; Braun 2012; Stein and Granot 2019). Pinto (2019) identified several SWEET transporters up-regulated in the roots of the Al-tolerant inbred line under Al-stress and the sucrose increment must be associated with these transporters playing a role in Al-stress in popcorn.

Sucrose synthase is involved in the conversion of sucrose into starch (Stein and Granot 2019). The degradation of starch in response to stress has been correlated with improved tolerance (Nagao et al. 2005; González-Cruz and Pastenes 2012). The content of starch decreased in the roots in the Al-tolerant inbred line, but increased in the shoots under Al-stress (Table 1). The starch degradation in roots can be correlated with the high sucrose content and the up-accumulation of sucrose synthase only in Al-tolerant inbred line due to the hydrolysis of previously stored starch in roots.

### 4.6. Glycolysis/gluconeogenesis

DAPs were detected in the glycolysis/gluconeogenesis pathway in all genotypes (Figure 3B). The putative phosphoglucomutase 1 (PGM1), cytoplasmatic (A0A1D6LQ82) was up-accumulated in the Al-sensitive and PGM1 cytoplasmatic (A0A1D6LQ83) was up-accumulated in both Al-tolerant genotypes. These proteins contribute to sucrose and cell wall components syntheses, and glucose phosphate partitioning (Malinova et al. 2014). The increase of glucose content in roots under Al-stress (Table 1), suggests that popcorn roots maintain the osmotic homeostasis of cells serving as an energetic resource in Al-stress conditions. For these reasons, we suggest that the glucose metabolism might be dependent on the glycolysis cycle in the Al-stress condition.

The proteins up-accumulated in both inbred lines could result in more glycolytically-generated ATP in roots to adapt to Al-stress, balancing the levels of available energy to prevent intracellular energy shortages. In rice roots, was found an accumulation increase of proteins in the glycolysis/gluconeogenesis pathway, suggesting that their play an important role in plant protection against Al^3+^ toxicity (Wang et al. 2014).

### 4.7. Carbohydrate metabolism

Organic acids (OAs) can prevent the entrance of Al in the root, chelating and immobilizing it on root surface (Kochian et al. 2004). In the Al-tolerant inbred line, pyruvate dehydrogenase E1 component alpha subunit (A0A1D6HJS3) and dihydrolipoamide acetyltransferase (A0A1D6QQE3) were up-accumulated. These proteins are involved in the pyruvate oxidation process. The pyruvate oxidation is an important connector that links the glycolysis to the rest of cellular respiration. These TCA enzymes were reported in the root proteome of cucumber (Du et al. 2010) and grapevine (Prinsi et al. 2020) under salt stress.

The three enzymes that form the core of the TCA cycle metabolon, citrate synthase (CS), isocitrate dehydrogenase (IDH), and malate dehydrogenase (MDH), were identified in our work with different accumulation patterns in at least one of the tested genotypes. In previous work, CS, MDH, and IDH were identified in the enhancement of Al tolerance in rice (Wang et al. 2014) and alfalfa (Sun et al. 2020). The up-accumulation of MDH (B4FG53) in the Al-sensitive inbred line may be associated with higher levels of malate in the roots (Table 1). However, the CS (B7ZWY9) was up-accumulated only in the Al-tolerant inbred line, and the fumarate hydratase and succinate dehydrogenase remained unchanged in the three genotypes (Suppl. Table S1). Overall, our results suggest that CS may play a crucial role in the chelation of Al and detoxification in the Al-tolerant inbred line, and malate and fumarate can help in the detoxification process, but are not the main players in the Al-tolerance in popcorn.

### 4.8. Integration of transcriptomic and proteomic data

Previously, Pinto (Pinto 2019) tracked the expression profile of these same inbred lines at the same time of Al exposure. With the integrative analysis, we are now able to detect that the response of Al tolerance in the Al-tolerant inbred line (11-133) is regulated mostly at the transcriptional level, but few components are shared at both levels (Figure 4A; Suppl. Table S2). The KEEG enrichment analysis showed that several important metabolic pathways in the Al tolerance response are regulated at the post-transcriptional level (Figure 4B). This integrative analysis provides useful information that may not have been deciphered on an individual platform (Haider and Pal 2013). Based on these results, the Al tolerance response provides insight into the transcription and translation dynamics of popcorn roots development and presents a complex regulatory network.

## 5. Conclusions

Protein expressions were associated with changes in protein synthesis, processing and degradation, oxidative stress, and carbohydrate metabolism. KEGG enrichment analysis indicated that sucrose metabolism was predominant activated in the Al-tolerant inbred line under Al-stress and proteomic analysis revealed that sucrose synthase should be an important player in the increasing of sucrose content and starch degradation to enhance popcorn tolerance to Al stress. Glycolysis/gluconeogenesis appeared to be a way to protect popcorn plants from Al^3+^ toxicity prevent intracellular energy shortage, as already visualized in other monocotyledons plants. Al stress induces malate and fumarate secretion preventing the Al^3+^ accumulation in the roots, and the regulation increased of citrate synthase suggests a key role of this enzyme in the detoxification process in the Al-tolerant inbred line. Finally, the Al-tolerance response is differentially regulated at transcriptional and post-transcriptional levels. These results highlight new players involved in the systemic plant response to Al^3+^ and help to understand the dynamic of plant-Al interaction in the root system and its contribution to Al-tolerance.

## Authors contributions

VBP designed the experiment, performed the proteomics analysis, and wrote the manuscript. VCA performed metabolites quantification, agronomical measurements, and statistical analysis. IAPL and WLA conducted metabolites quantification. EMV performed the proteomics analysis. WLA, VS, and JMSV provided guidance for the study, participated in the design and reviewed the manuscript. JMSV provided funds for the study. All authors read and approved the manuscript.

## Acknowledgments

We thank the National Council for Scientific and Technological Development (CNPq), the Brazilian Federal Agency for Support and Evaluation of Graduate Education (Capes; Finance Code 001), and the Foundation for Research Support of Minas Gerais State (Fapemig) for financial support.

## Supplementary material

**Supplementary Figure 1** Sucrose content of roots and shoots during 24; 72; and 168 hours under Al-stress. Black bars indicate control condition and gray bars indicate Al treatment. Average followed by the same lower-case letters (time) and capital letters on the columns (environment) do not differ significantly by the Tukey test (P<0.05), (n=10).

**Supplementary Table 1** Comparative proteomic analysis of popcorn roots under 72h of Al-stress.

**Supplementary Table 2** Proteins and transcripts shared between proteomics and transcriptomics analyzes in Al-tolerant inbred line under Al-stress.

