## Supplementary figures and images for "Deciphering the major metabolic pathways associated with aluminum tolerance in popcorn roots using label-free quantitative proteomics"

### Suppl. Figure S1

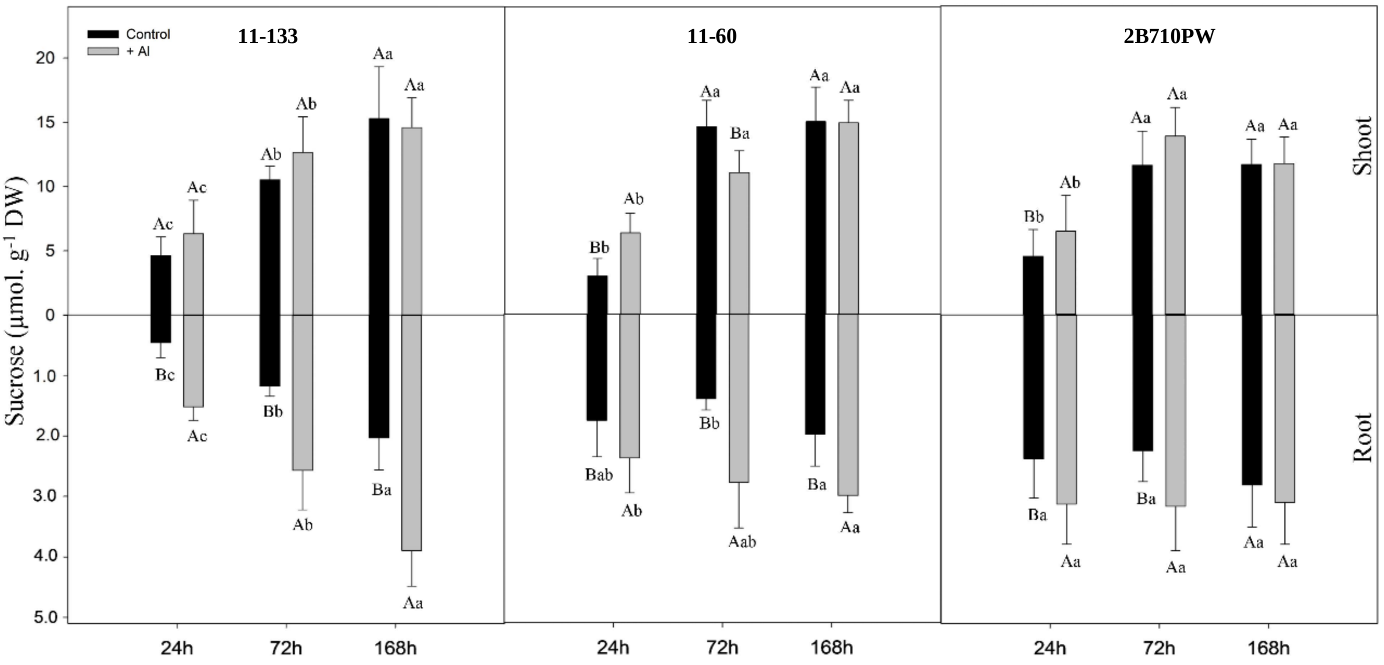
